# Detection of DNA replication errors and 8-oxo-dGTP-mediated mutations in *E. coli* by Duplex DNA Sequencing

**DOI:** 10.1101/2022.10.21.513255

**Authors:** Niketa Bhawsinghka, Adam Burkholder, Roel M. Schaaper

## Abstract

Mutation is a phenomenon inescapable for all life-forms, including bacteria. While bacterial mutation rates are generally low due to the operation of error-avoidance systems, sometimes they are elevated by many orders of magnitude. Such a state, known as a hypermutable state, can result from exposure to stress or to harmful environments. Studies of bacterial mutation frequencies and analysis of the precise types of mutations can provide insights into the mechanisms by which mutations occur and the possible involvement of error-avoidance pathways. Several approaches have been used for this, like reporter assays involving non-essential genes or mutation accumulation over multiple generations. However, these approaches give an indirect estimation, and a more direct approach for determining mutations is desirable. With the recent development of a DNA sequencing technique known as Duplex Sequencing, it is possible to detect rare variants in a population at a frequency of 1 in 10^7^ base pairs or less. Here, we have applied Duplex Sequencing to study spontaneous mutations in *E. coli*. We also investigated the production of replication errors by using a mismatch-repair defective (*mutL*) strain as well as oxidative-stress associated mutations using a *mutT*-defective strain. For DNA from a wild-type strain we obtained mutant frequencies in the range of 10^-7^ to 10^-8^ depending on the specific base-pair substitution, but we argue that these mutants merely represent a background of the system, rather than mutations that occurred *in vivo*. In contrast, bona-fide *in vivo* mutations were identified for DNA from both the *mutL* and *mutT* strains, as indicated by specific increases in base substitutions that are fully consistent with their established *in vivo* roles. Notably, the data reproduce the specific context effects of *in vivo* mutations as well as the leading vs. lagging strand bias among DNA replication errors.

## 1. Introduction

*E. coli* has been a model organism for genetic analysis for many decades [1], and it is still widely used to gain insights into the fundamental mechanisms by which cells maintain genome integrity. It is also a popular expression platform for the production of recombinant proteins, some of which have found their way into therapeutics. Being such an important genetic work-horse, there needs to be a clear picture of the mutations occurring in its DNA and the mechanisms by which they occur or are prevented. Under normal circumstances, mutation rates in *E. coli* are low (10^-9^ to 10^-10^) [2] due to the operation of a number of pathways contributing to the integrity of the genome. Base selection, exonucleolytic proofreading and post-replicative DNA mismatch repair operate serially to ensure high intrinsic fidelity of DNA replication [3], although some errors may escape this scrutiny and will end up as mutations. DNA damage from a variety of endogenous and exogenous sources, if left unrepaired, is also expected to be a significant contributor to spontaneous mutations. Transcription, replication-transcription conflicts and movement of transposable sequences are further potential contributors to sources of mutations [4].

Several approaches have been used to estimate the level of spontaneous mutations in this organism. Most of these studies, including many from our laboratory, have used long-established methods such as fluctuation assays and clone-based sequencing that rely on the selection of detectable phenotypes [5, 6]. More recently, studies using whole-genome sequencing after multiple generations of mutation accumulation have been reported [7]. Estimation of *E. coli* mutation rates have also been reported using comparative genomics [8]. The frequencies and mutational spectra from these studies tend to agree at some level, but also show significant differences, as discussed in [9]. Techniques using mutant selection represent indirect measurements that require many assumptions and extrapolations [9], while mutation accumulation methods although less subject to selection issues are, instead, quite laborious and time consuming. A detection of mutations directly in DNA without the need for selection or extensive genetic manipulations may have significant advantages in this respect.

Duplex DNA Sequencing is a recently developed technique, which can detect mutations directly in isolated DNA at potentially very low frequencies [10]. The approach relies on the tagging and sequencing each of the two strands of unique DNA duplex fragments. Bona-fide mutations existing in double-stranded DNA are identified by their presence at the same position in the two complementary strands and therefore can be used readily for detecting mutations in heterogeneous populations. The technique has been used to detect mutations in mitochondria and within tumors, and mutant frequencies in the range of 10^-5^ to 10^-7^ have been reported [11, 12] which is a significant improvement over the standard next generation sequencing (NGS) error rate (10^-2^ to 10^-3^) [13].

In the present study, we applied the Duplex Sequencing methodology to investigate mutations occurring in the bacterium *E. coli* as a continuation of the multiple studies of mutagenesis in this organism by traditional methods. Specifically, we used *E. coli* BL21-AI, a member of the B-lineage of *E. coli* [14], which is 98.8 % identical to the more traditionally used *E. coli* K-12. The B strains have been used most prominently for studies and applications of protein expression and overproduction [15]. Specifically, BL21-AI is a derivative that is used in conjunction with T7-based expression vectors to achieve more controlled expression of harmful or toxic proteins due to a tighter control of basal expression levels [14]. Tighter control is achieved by the presence of an ara-T7pol cassette in its chromosome, in which the expression of the T7 RNA polymerase gene is controlled by the tight arabinose-inducible (and glucose-repressible) P_BAD_ promoter. DNA sequencing of the BL21-AI genome has shown the chromosomal sequence to be essentially identical to that of the parental *E. coli* B, except for a 4-kb insertion of the araC-T7 pol cassette (introduced from an *E. coli* K-12 strain by P1 transduction) [14]. As shown in this work, the mutability of BL21-AI as measured by the appearance of rifampicin-resistant mutations is low and comparable to frequencies obtained for the K-12 strains, justifying its use in the present investigation.

The DNA duplex sequencing was executed on a ~10.3-kb region of the BL21-AI genome that in addition to normal *E. coli* B sequences also included the genes involved in recombinant protein expression (*ara* operon, RNAP gene, *tetB*). The 10-kb region is shown in Fig. 1. We first used the wild-type BL21-AI strain to investigate any background frequency that could be detected using Duplex Sequencing. We then expanded our investigation by creating two genetic variants of the strain: (a) a *mutL* derivative to potentially detect DNA replication errors and (b) a *mutT* derivative to identify mutations resulting from oxidative stress. Both strains are well established mutators that show elevated mutation rates with a characteristic mutational fingerprint that relates to their underlying error prevention/production systems. Our results show that the Duplex Sequencing approach is indeed capable of revealing the specific classes of base-substitution mutations produced in both these strains, and that these mutations are readily detected above the background of the system. Also, we also address issues with the present background of Duplex Sequencing, which may help in further refinement of this technique and expansion of its applicability.

**Fig. 1:**
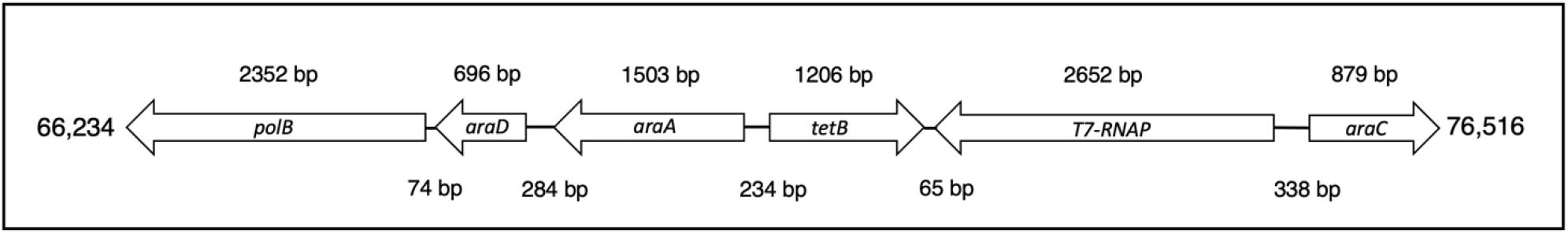
Representation of 10.3-kb chromosomal region (genomic coordinates 66,234-76,516) of *E. coli* BL21-AI [14] used for Duplex sequencing. The region has normal *E. coli* B genes *(polB)* as well as genes involved in protein-expression (ara, *tetB*, T7-RNAP). Numbers in the upper panel depict the length of coding genes and those in the bottom panel the length of the intergenic regions.

## 2. Materials and methods

### 2.1. Bacterial strains and Media

*E. coli* strain BL21-AI [14] and its derivatives were used for all experiments, and genetic deficiencies were introduced into this strain by P1 transduction (REF, Miller). The mismatch-repair-defective BL21-AI *mutL* derivative NR18632, in which the *mutL* gene is inactivated by insertion of transposon Tn*5*, was constructed by transferring the *mutL*::Tn*5* allele (conferring kanamycin resistance) from donor strain NR9559 [16] using phage P1 *cml crl100* (Miller 1972) as described [17]. The *mutT*-deficient derivative NR18661, in which the *mutT* gene is replaced by the Δ*mutT*::*cat* allele (conferring chloramphenicol resistance), was created by first generating the Δ*mutT*::*cat* allele in strain BW25113/pKD46 using the PCR-based method of Datsenko and Wanner [17]. The source of the *cat* allele was plasmid pKD3 and primers used for the PCR reaction were: 5’-GTTGAAGTAAAAGGCGCAGGATTCTGCGCCTTTTTTATAGGTTTAAGACAGTGTAGGCTGGAG CTGCTTC-3’ (forward) and 5’-ATCAGGCATCGTGTGCGAATGTCGGATGCGGCGAAAACGCCTTATCTG ACCATATGAATATCCTCCTTAG-3’ (reverse). Subsequently, the Δ*mutT*::*cat* allele was transferred to BL21AI using P1 transduction with P1*virA*. Mutator phenotypes were confirmed by testing for elevated frequency of rifampicin-resistant mutants (see below). LB medium was used throughout.

### 2.2. Mutant frequency determinations using rifampicin resistance

Determination of mutant frequencies was done as described [18]. Briefly, for each strain a total of 30 independent overnight cultures were grown at 37°C, each started from a single colony. The number of mutants was determined by plating 100 μl of the overnight cultures on LB plates containing 100 μg/ml rifampicin followed by incubation at 37°C for ~20 hrs. The total number of cells was obtained by plating 100 μl of a 10^-6^ dilution of the cultures on LB plates. For mutant frequencies, the median number of mutants obtained from the 30 cultures was divided by the average CFU.

### 2.3. Genomic DNA Isolation

Genomic DNA (gDNA) was isolated from 4 ml of overnight cultures using the Qiagen Genomic-tip 100/G as per manufacturer’s instructions. The quality of the extracted DNA was evaluated by agarose gel electrophoresis and measurement of the A260/A280 ratio. DNA quantification was done using Qubit dsDNA HS Kit (Invitrogen).

### 2.4. DNA library preparation and sequencing

100 ng of gDNA was sheared using a Covaris E220 Focused-Ultrasonicator in microtubes-50 to yield an average fragment size of 300 bp. DNA libraries were prepared using TwinStrand Duplex Sequencing Universal Kits (Twinstrand Biosciences, Seattle, WA) following manufacturer’s instructions with some modifications. Briefly, the procedures involved (i) end repair and A-tailing of the DNA fragments, (ii) ligation of duplex adaptors at both fragment ends, (iii) a “conditioning” reaction using a mix of DNA repair enzymes (Twinstrand) to eliminate common forms of DNA damage, (iv) ten rounds of “indexing” PCR amplification (PCR-1) with indexing primers, (v) two rounds of enrichment (by hybridization) for fragments corresponding to a 10.3-kb genomic region (coordinates 66,234 – 76,516) of the BL21AI genome [14] using custom-designed biotinylated ssDNA probes (Agilent SureSelect) and streptavidin-bead purification. Two additional rounds of PCR amplification were included: PCR-2 (16 cycles) was performed between the two enrichment steps, and PCR-3 was performed following the final enrichment (8 cycles or more, as needed to obtain sufficient product to allow Qubit quantitation). Bead purification of the DNA was performed between all consecutive steps. The final libraries were quantitated by the Qubit dsDNA HS method and analyzed on an Agilent 2100 Bioanalyzer. Indexed libraries were pooled, and DNA sequencing was performed on Illumina NextSeq 500 or NovaSeq 6000 sequencers using 150-bp paired-end reads. A requirement for success in these experiments is the use of an appropriate (and limited) amount of starting DNA fragments in the PCR-1 amplification [13]. This amount ultimately determines the so-called “family size”, which represents the number of times unique individual DNA fragments, including those having a mutation, are found in the final Illumina sequence output. This number has been recommended to be six or greater before a mutation is to be accepted as a valid mutation originally present in both DNA strands [13]. In our experiments, we typically used 0.8 to 2 ng of adapter-ligated library DNA for PCR-1, which produced family sizes in the range of 10-50. The total number of Illumina reads was in the range of 30-100 million, while overall duplex depth (the number of consensus reads for each base pair in the target) was in the range of 10,000. At this depth, a single mutation noted in the 10^4^-bp target would represent a frequency per bp of 10^-8^.

### 2.5. Bioinformatic analysis

The Twinstrand duplex adapters contain 8-nt duplex barcodes of degenerate DNA sequence, and these barcodes are used to identify read families that correspond to unique, original double-stranded DNA fragments. Using the FastqToBam module of the fgbio package, version 1.5.1 (http://fulcrumgenomics.github.io/fgbio/tools/latest/FastqToBam), the 8-nt barcode and a 1-nt spacer were removed from the 5’ ends of each read and the paired barcode sequences were included as a metadata tag attributed to both reads in the output BAM file. This BAM was subsequently converted to FASTQ format and aligned to the E. coli strain BL21-AI genomic sequence [14] using BWA-MEM 0.7.15-r1140 [19] with a reduced clipping penalty of 1 (-L1,1). The aligned reads for each sample were merged with the existing barcode metadata using the Picard Tools 2.26.11 module, MergeBamAlignment (https://broadinstitute.github.io/picard/). Read families were then generated from the barcode tags using fgbio module GroupReadsByUmi (http://fulcrumgenomics.github.io/fgbio/tools/latest/GroupReadsByUmi) employing the paired UMI grouping strategy (--strategy=paired). Quality control (QC) information was then extracted from the resulting BAM file using fgbio module CollectDuplexSeqMetrics (http://fulcrumgenomics.github.io/fgbio/tools/latest/CollectDuplexSeqMetrics) and the resulting outputs were examined for bias that might affect consensus calling or identification of mutations in each sample. Duplex consensus reads were then generated from those previously grouped by barcode using fgbio module CallDuplexConsensusReads (http://fulcrumgenomics.github.io/fgbio/tools/latest/CallDuplexConsensusReads). This process included trimming of low-quality 3’ ends, masking bases having individual quality scores below 20 with ‘N’, and enforcing a minimum family size of 6 with a minimum of 3 read pairs from each strand, but otherwise utilized default parameters (--trim --min-reads 6 3 3--min-input-base-quality 20). Filtering of duplex reads was then performed using fgbio FilterConsensusReads (http://fulcrumgenomics.github.io/fgbio/tools/latest/FilterConsensusReads), masking bases derived from fewer than 6 read pairs or fewer than 3 supporting each strand, those with a consensus quality score less than 40, those with an error rate greater than 0.1, and those where the single-strand consensus bases did not agree (-M 6 3 3-N 40-E 0.1-s). Duplex read pairs with an overall error rate greater than 0.3 were also excluded at this stage (-e 0.3). Finally, filtered and masked duplex reads were aligned to the BL21-AI genomic sequence using bwa mem (-L 1,1) and the aligned pairs were hard-clipped using fgbio module ClipBam (http://fulcrumgenomics.github.io/fgbio/tools/latest/ClipBam) to remove 7 bases from the 5’ end of each read as well as any overlapping 3’ sequence to reduce false positives and avoid double-counting mutations falling in the overlapping ends (-c Hard--read-one-five-prime 7-read-two-five-prime 7--clip-overlapping-reads). Mutations were then identified within the target region of positions 66234-76516 using VarDictJava 1.8.2 [20], with parameters tuned to report those supported by a single read or more (-f 0 -r 1 -P 0 -p -m 9999). The total count of bases aligned to each position within the target was determined separately by reference base identity (A/C/G/T) using a combination of samtools 1.3.1 and short custom scripts. These values were combined with the output produced by VarDict to determine mutation counts and frequencies for each type of base substitution *(e.g*., A→T, G→A, C→G). To facilitate further QC and data examination, bedGraph formatted tracks displaying duplex depth for each target position as well as the fraction of masked (N) bases per position were generated from the aligned and clipped duplex reads obtained from the samtools mpileup (http://www.htslib.org/doc/samtools-mpileup) output. Additionally, duplex read pairs containing non-reference bases were identified using samtools calmd (http://www.htslib.org/doc/samtools-calmd), and with the aid of custom scripts and Picard tools MergeBamAlignment, a BAM file was generated for each sample that contained all observed alternate alleles as well as all consensus calling metadata.

## 3. Results

### 3.1. Duplex Sequencing on wild-type *E. coli* DNA

Even after the operation of multiple error-avoidance mechanisms in *E. coli*, as in the wild-type (WT) strain, mutations do occur at a low frequency. To investigate whether Duplex Sequencing method can be used to obtain insight into the spontaneous (or background) mutations we applied the method to the wild-type BL21-AI strain. Five independent colonies of the strain were inoculated to create overnight cultures. Genomic DNA was extracted and duplex libraries prepared for DNA sequencing using manufacturer recommended procedures (Twinstrand Biosciences, see Methods). The libraries were enriched (see Methods) for a ~10.3 kb region of the BL21-AI genome, and the sequencing results we report are for this 10.3 kb target (Fig. 1). The total number of reads per library varied from 30-100 million, while the median duplex depth (the number of times each given base-pair in the target was read) was 11,650, with uniform representation across the target. The ‘family size’, which represents the number of times a PCR copy of a given unique chromosomal DNA fragment is represented in a sequencing run (based on identical tag combinations) ranged from 10-50. This sufficiently high value allowed us to satisfy the duplex sequencing criteria (see Methods) for accepting potential mutations as original mutations present in both DNA strands (but see Discussion).

Over the five independent experiments, a total of 401 mutants (refer Fig.4) were obtained that passed the bioinformatics criteria. The mutant frequency for the different libraries ranged from 1.5 x 10^-7^ (19 mutations out of 123,211,935 duplex bases) to 4.2 x 10^-7^ per bp (41 mutations out of 97,518,372 duplex bases) with an average mutant frequency for the wild type of 2.7 x 10^-7^ per bp. Importantly, as shown in in Table 1, the calculated frequencies for each of the six different types of base-pair substitutions varies considerably. Mutation frequency at A·T sites are generally significantly lower (~10 fold) than at GC sites. Among A·T substitutions, A·T→T·A has the lowest mutation frequency (in most libraries no mutations were detected among several million duplex bases). Substitutions at G·C sites were higher, notably G·C→C·G and G·C→T·A transversions, which exceeded the combined transitions by 10-fold (Fig. 2a). Although a few short indels were also detected, we focused our analysis on the single base-pair substitutions.

**Table 1.** Mutant frequencies obtained by Duplex Sequencing for the wild-type strain. Mean and standard error of mean (SEM) are represented (n=5). Substitutions at G·C sites are significantly higher than at A·T sites (χ^2^= 54.4, P= 4.2 x 10^-11^).

| Type of mutation | Mean mutation frequency | SEM |
| --- | --- | --- |
| A·T substitutions | $8.9 \times 10^{-8}$ | $2.7 \times 10^{-8}$ |
| A·T → G·C | $3.7 \times 10^{-8}$ | $1.3 \times 10^{-8}$ |
| A·T → T·A | $3.8 \times 10^{-8}$ | $1.7 \times 10^{-8}$ |
| A·T → C·G | $1.3 \times 10^{-8}$ | $3.5 \times 10^{-9}$ |
| G·C substitutions | $4.3 \times 10^{-7}$ | $1.2 \times 10^{-7}$ |
| G·C → A·T | $1.0 \times 10^{-7}$ | $1.1 \times 10^{-8}$ |
| G·C → C·G | $1.5 \times 10^{-7}$ | $5.9 \times 10^{-8}$ |
| G·C → T·A | $1.7 \times 10^{-7}$ | $5.4 \times 10^{-8}$ |

**Fig. 2:**
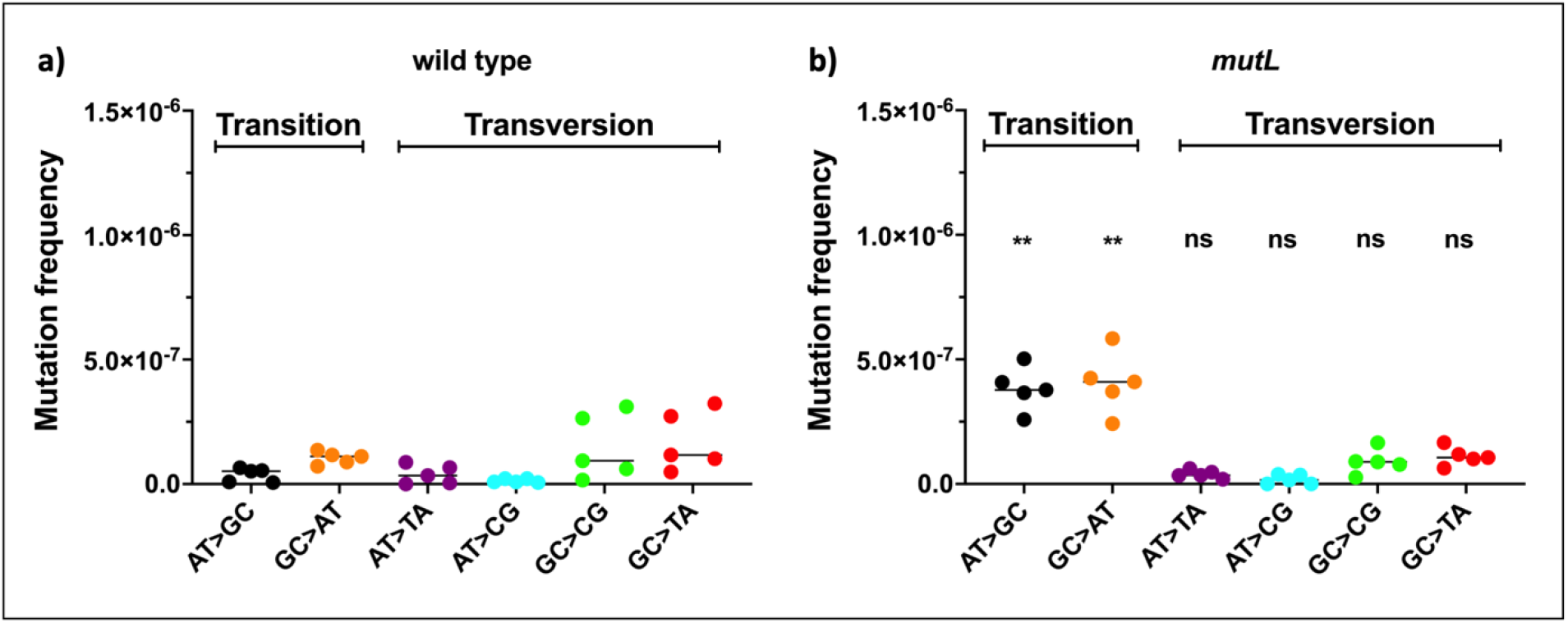
Mutant frequencies and specificity of base-pair substitutions in (a) the wild-type (WT) and (b) the *mutL* strain. The results of five independent experiments are shown along with median. Mann-Whitney U test was performed to determine the significance of changes in *mutL* compared to WT (each type of mutation in *mutL* has been compared to the respective type in WT). The changes for A·T→G·C (P = 0.0079, **) and G·C→A·T (P = 0.0079, **) are statistically significant, whereas the changes for the others are non-significant (ns).

While these frequencies, in the range of 10^-7^ to 10^-9^, are quite low and testify to the sensitivity of the Duplex Sequencing approach, the frequencies are still significantly higher than expected based on estimations of *in vivo* mutation rates obtained from genetic experiments. As outlined in the Discussion, various approaches have indicated that the mutation rate per base pair in *E. coli* is as low as 10^-10^ (10^-9^ – 10^-11^) [3, 7, 18]. Thus, we conclude that the data obtained here for the wild-type strain do not reflect actual *in vivo* mutations, but represent a background of the system. The difference in frequencies for substitution at A·T and G·C base pairs (Table 1) also supports this conclusion, as transitions and transversions contribute roughly equally to the spontaneous *in vivo* background [3, 21]. The issue of background and its possible sources is further addressed in the Discussion. Thus, although Duplex Sequencing is currently the most sensitive technique to detect mutations directly in DNA, it is not currently sensitive enough to determine the true mutation frequency in wild-type *E. coli*. Nevertheless, the results from this experiment have the potential to be exploited for understanding the contributing factors to the present background level. In the following section, we describe additional experiments where we attempt to elevate the *in vivo* mutation rate by introducing genetic mutations in important mutation avoidance systems, which may allow the mutation frequencies to rise above the background of the system.

### 3.2. Detection of primary replication errors in a *mutL* strain

The *in vivo* mutation rate in *E. coli* is low due to the action of several mutation prevention systems. One such system is the DNA mismatch repair system (MMR), which depends on the concerted action of the *mutH, mutL*, and *mutS* gene products [22]. Via the MMR system, DNA polymerase errors (*i.e*., polymerase-introduced misinsertions that escape removal by the associated exonucleolytic proofreading) are scrutinized post-replicatively and corrected through repair synthesis on the newly synthesized DNA strand. The MMR system is highly efficient as judged by the 100-to 1,000-fold increase in the *in vivo* mutation rate when it is disabled by a defect in either of the *mutH, mutL* or *mutS* genes [23]. This increase is primarily of A·T→G·C and G·C→A·T transitions, as these are the replication errors specifically suppressed by the MMR system [23]. Thus, we asked whether duplex sequencing might be able to reveal such uncorrected replication errors under conditions of disabled MMR. For this we constructed a *mutL*-defective BL21-AI derivative (see Methods) and confirmed the mutator phenotype using the conventional rifampicin-resistance system [24]. As shown in Table 2, the *mutL* strain showed a near 200-fold increase in the Rif^r^ mutant frequency relative to the wild-type strain.

**Table 2.**
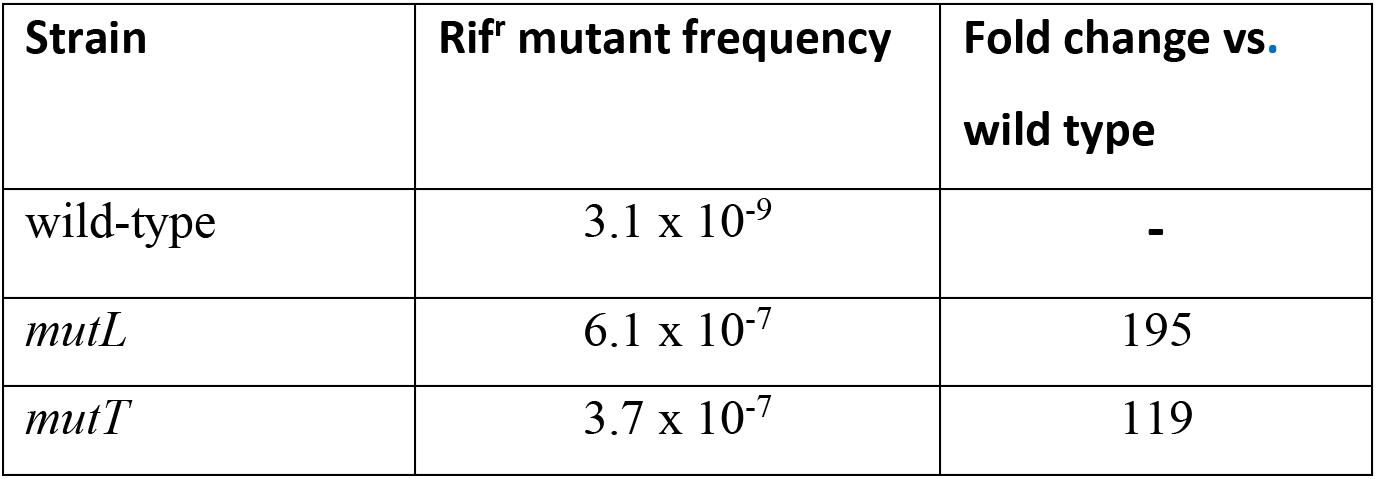
Mutant frequencies obtained using the Rif^r^ fluctuation assay. The means from four independent experiments are shown. Each independent experiment was done using 30 overnight cultures.

We then performed Duplex Sequencing on five independent libraries prepared from the *mutL* strain. The average mutation frequency obtained was 5.3 x 10^-7^, which represents a roughly 2-fold increase over the frequency of the wild-type (2.7 x 10^-7^). Interestingly, as revealed in Fig. 2, the individual base substitution classes are affected differentially: while in the wild-type the transversions dominated, the opposite is true for the *mutL* strain, where A·T→G·C and G·C→A·T transitions dominate. The data obtained (Fig. 2) show that the frequency of transversions remains essentially unchanged between the two strains, but there is a specific frequency increase for the two transitions, about 10-fold for A·T→G·C and about 4-fold for G·C→A·T (Fig.2b). We conclude from these results that we have been successful in using Duplex sequencing to reveal the contribution of primary replication errors. The specific increase for the transitions is fully consistent with the established *in vivo* mutational specificity of MMR-defective strains [7, 23], where A·T→G·C and G·C→A·T transitions outnumber by some two orders of magnitude, and this has been interpreted to indicate that the primary function of the MMR system is to prevent the occurrence of transitions, which constitute the bulk of the uncorrected replication errors [18]. Of course, we note that the observed 10-to 4-fold increase for transitions is less than observed *in vivo* experiments for *mutL*-defective strains (Table 2 and [3]), but this is a direct result of the higher background in the duplex sequencing experiment (Table 1).

### 3.3. Detection of mutations generated due to 8-oxo-dG misincorporation in replicating DNA

The success with detecting primary replication errors in the *mutL* background using the Duplex Sequencing method led us to employ another *E. coli* mutator strain, *mutT*. This established mutator is known to produce a strong and specific increase for A·T→C·G transversions [25]. The source for these mutations is the presence of 8-oxo-dGTP in the dNTP precursor pool, which is a frequent oxidative stress product resulting from the oxidation of dGTP. The normal level of 8-oxo-dGTP is kept very low due to the action of the *mutT* gene product. MutT is an 8-oxo-dGTP pyrophosphohydrolase that inactivates 8-oxo-dGTP by converting it to 8-oxodGMP and pyrophosphate. If not removed, 8-oxo-dGTP is a powerful mutagen, as it is readily incorporated by the DNA polymerase opposite template adenines [26]. Once in the DNA, 8-oxo-dG may pair with dCTP in the next round of replication, resulting in an A·T→C·G transversion. This is a powerful mutational mechanism as evidenced by the 1,000-to 10,000-fold increase in A·T→C·G transversions in a *mutT*-defective background as measured by *in vivo* systems employing mutant selection [25, 27].

We constructed a *mutT*-deficient derivative of BL21-AI (see Methods), and measurement of the Rif^r^ frequency confirmed the mutator phenotype (see Table 2). DNA libraries for Duplex Sequencing were prepared from six independent colonies. The average mutant frequency was 6.6 x 10^-7^, an about 2.5-fold overall increase (6.6 x 10^-7^ vs. 2.7 x 10^-7^) over wild type. However, and importantly, the detailed results show that the only significant spectral change is a strong increase (nearly 40-fold), in A·T→C·G transversions (Fig. 3). In fact, among total of 763 mutations detected in *mutT* DNA, randomly distributed over the entire target (Fig. 4), 433 were A·T→C·G. We conclude that we have successfully applied the Duplex sequencing method for the detection of 8-oxo-dGTP-mediated mutations under conditions where the levels of the mutagenic intermediate are elevated.

**Fig. 3:**
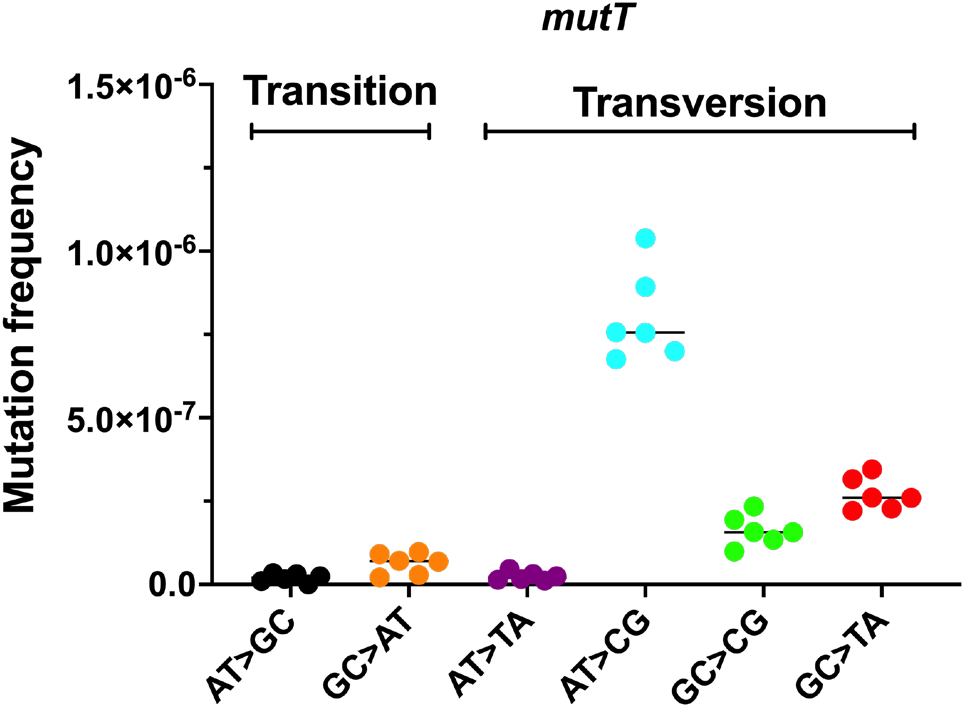
Mutant frequencies and base-pair substitution specificity in a *mutT*-deficient strain revealed by Duplex sequencing. Six independent measurements along with median are shown. The change in frequency of A·T→C·G is statistically significant (P = 0.0043, **) as compared to WT (Fig. 2a) using Mann-Whitney U test; changes for the other substitutions are not significant.

**Fig. 4:**
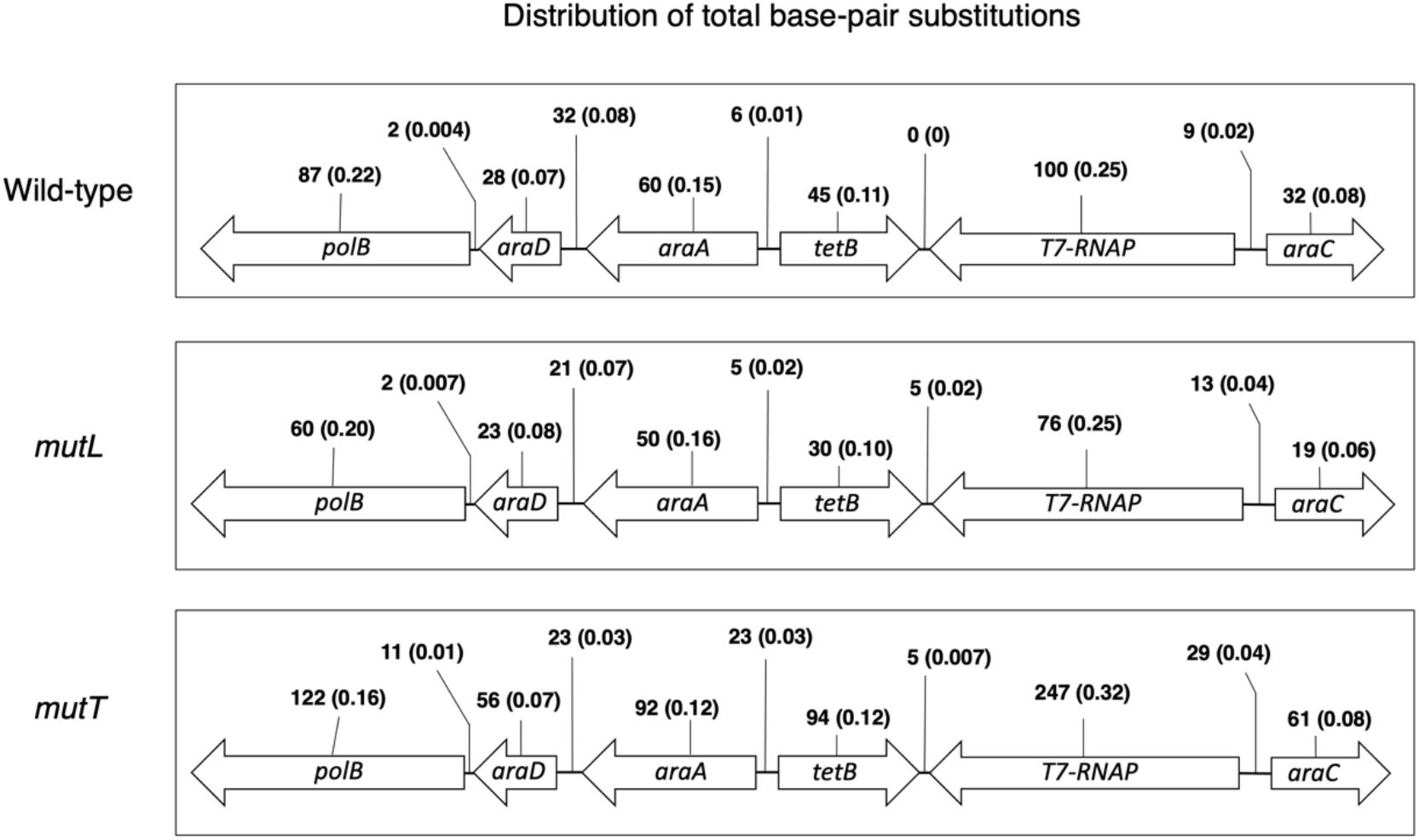
Distribution of the base-pair substitution mutations obtained by Duplex DNA sequencing in three different strains as indicated on left. The cumulative number of mutations for each strain is listed above the genes and the intergenic regions along with the fractions (in parentheses) relative to the total number of mutations (401 for wild type, 304 for *mutL*, 763 for *mutT*).

## 4. Discussion

DNA Duplex Sequencing is an important new technique that allows for the direct detection of mutations in isolated DNA. The approach avoids the need for phenotypic selection of mutant cells or any long-term mutation accumulation efforts, both of which must be followed by the sequencing of individual clones. Importantly, in the duplex method unique tagging of both strands of ds DNA fragments greatly facilitates the scoring of mutations present at low frequencies in heterogeneous populations. Here, we have investigated Duplex Sequencing in the bacterium *E. coli* with an eye towards understanding the meaningful application of the approach to questions of interest regarding bacterial mutagenesis. Overall, the results show that duplex sequencing is a sensitive method for mutation detection in bacterial DNA, yielding an observed average mutant frequency for wild-type DNA of 2.7 x 10^-7^ per base-pair. It is important to note that this frequency depends strongly on the type of base substitution, as it ranges from a low of 2.1 x 10^-8^ for mutations at A·T base-pairs to a high of 1.7 x 10^-7^ at G·C base-pairs. Our data are solid, based on a total of 401 detected mutations (Table S1) distributed randomly across the 10,282-bp long chromosomal target (Fig. 4). The unbiased distribution of mutations across the coding and non-coding regions shows that the 10.3-kb region chosen for mutation scoring reflects a non-biased representation of the entire genomic DNA, and conclusions should be extendable to the chromosome at large.

At the same time, we conclude that the frequency numbers are too high to represent actual *in vivo* mutation rates, most obviously for the G·C→T·A and G·C→C·G transversions which appear at frequencies exceeding 10^-7^ (Table 1). Several *in vivo* studies have revealed that the *in vivo* mutation rate is very low, benefitting from the sequential and parallel operation of several mutation avoidance systems [3, 9], resulting in a calculated *in vivo* mutation rate of 1-2 x 10^-10^ [3, 27]. Thus, we conclude that the observed frequencies in the duplex sequencing experiment reflect a background of the system. A likely source of background mutations is the introduction of DNA damage in the duplex library preparation, including the steps of DNA isolation and shearing and including the subsequent amplification steps. Particularly, the likelihood of introducing oxidative and heat-induced damage during the ultrasonicator-mediated DNA shearing has been advanced [28, 29]. Particularly guanine bases are prone to being oxidatively damaged, creating products like 8-oxoguanine and further oxidation products, of which several have been identified [30, 31]. These products have been shown to be highly mutagenic, yielding G·C→T·A and G·C→C·G transversion mutations [32], and this specificity is consistent with the high frequencies for these events seen in our study (Figs. 2 and 3), including their sequence specificity (Tables S2 and S3). The data show a preference for the G·C→T·A events to occur at 5’-G or 5’-C (5’-G/C **<u>G</u>**) and 3’-T (**<u>G</u>** T-3’) (where **<u>G</u>** is a template base undergoing the change) and a bias for 5’-T (5’-T**<u>G</u>**) and 3’-A (**<u>G</u>**A-3’) for the G·C→C·G transversions. DNA library preparation is obviously an area where improvements can be made, for example using methods that avoid sonication. It should also be noted that DNA lesions introduced during library production are expected to be present generally in only one of the two DNA strands. Therefore, as the duplex sequencing method is designed to detect only mutations present in both DNA strands of the starting DNA, such single-strand events should not lead to detectable mutations using this method. We assume that DNA strand loss during some of the early amplification steps, even if generally infrequent, is one of the reasons by which heteroduplex molecules can be recovered as seemingly bona-fide duplex mutations. This is another potential area where improvement might be made.

### Analysis of DNA replication errors

Despite the background issues discussed above, important and relevant results were obtained with chromosomal DNA obtained from a mismatch-repair defective (*mutL*) strain. Based on a total of 304 retrieved mutations (Fig. 4 and Table S1), we observed a specific increase of A·T→G·C and G·C→A·T transitions of ~10- and ~4-fold above their respective backgrounds (Fig. 2). This specificity is in full agreement with the mutational spectra observed for mismatch-repair defective *E. coli* strains using phenotypic assays [23] or mutation accumulation experiments [7], which reveal a large predominance of these two types of transitions. This strong transition preference reflects the nature of primary replication errors, which are predominantly of the transition type. Note that our frequencies of A·T→G·C and G·C→A·T correspond well to their estimated *in vivo* rates of around 10^-7^ [3, 7, 18].

Inspection of the DNA sequence contexts of the observed A·T→G·C and G·C→A·T events additionally confirms their correspondence with *in vivo* mutations. For example, for A·T→G·C (normalizing to errors at template <u>T</u> residues) there is a significant preference for a 5’-G (58 %) or a 3’-A (34 %) with a full 20 % occurring in the context 5’-G**<u>T</u>**A-3’. For the G·C→A·T (normalized to template <u>G</u> residues) 39 % have a 5’-C and 47 % have a 3’-C, with 21 % occurring in the sequence context 5’-C**<u>G</u>**C-3’. These neighboring base preferences correspond well to those derived from *in vivo* mutation accumulation studies [7] or *in vivo lacI assays* [23], further validating the nature of these mutations as (uncorrected) *in vivo* replication errors (Table S4 and S5).

Most importantly, our *mutL* results confirm a distinct strand bias in the occurrence of the transition mutations. Such strand biases can be revealed when making assumptions about the mispairings underlying the two transitions. A·T→G·C can result in principle from replication errors at template <u>T</u> sites (<u>T</u>·G errors) or template <u>A</u> sites (<u>A</u>·C errors). However, <u>T</u>·G errors are generally assumed significantly more frequent than <u>A</u>·C errors [18, 33], and this bias can be used to decipher the strands in which the A·T→G·C originated. Our data show that with respect to the BL21AI reference (+) strand, among 99 A·T→G·C events the majority (81) are observed as A→G while only 18 are T→C (Fig. 5). It follows then that the majority arises as T·G in the non-reference (-) strand (χ^2^= 40.5, P= 2 x 10^-10^). In view of the location of our 10.3 kb target (coordinate 66,234 - 76,516) relative to the origin of replication (OriC) [14] (Fig.6), this (-) strand is copied by the leading-strand replication machinery. Thus, our results suggest that most errors are associated with leading-strand replication and that lagging-strand replication has higher fidelity. This is fully consistent with earlier reports [18, 33] that ascribed a higher fidelity to the lagging strand of replication. The same or similar result is obtained for the G·C→A·T errors, which arise predominantly due to errors at template G sites (G·T errors). Out of 119 observed G·C→A·T substitutions in the reference (+) strand, 66 % (79 out of 119) are observed as C →T and the rest (34%) as G→A changes (χ^2^= 10.7, P= 0.001) (Fig. 5). We conclude, again, that most of these errors arise out of leading-strand synthesis (<u>G</u>·T errors). These results are a powerful outcome validating the Duplex sequencing approach as a means to study issues of replication fidelity in bacteria and, presumably, other organisms.

**Fig. 5:**
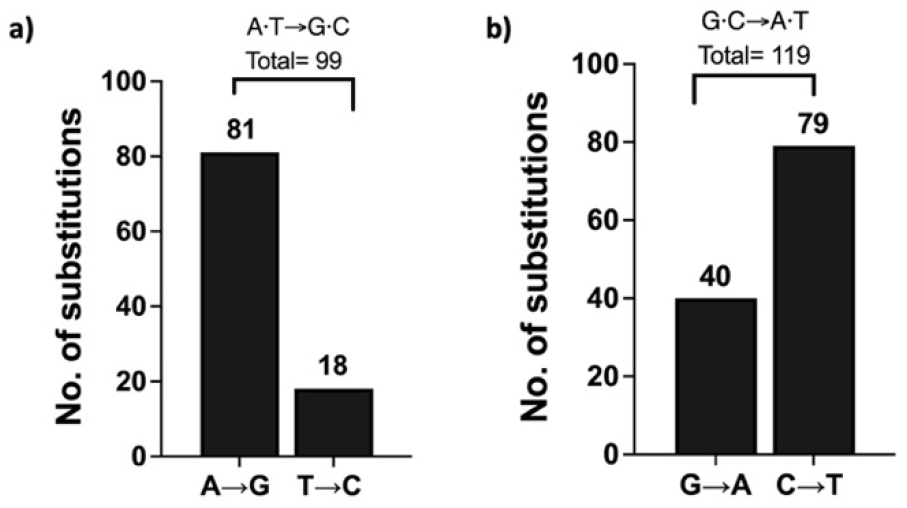
Strand Bias among replication errors in a *mutL* strain.

**Fig. 6:**
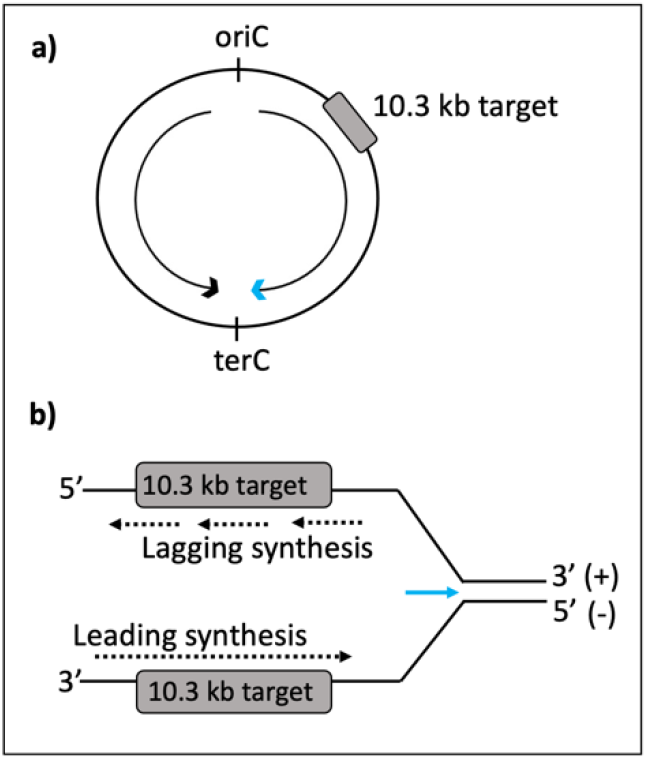
Location of the target sequence (grey boxes) on the BL21AI chromosome with respect to origin of replication (oriC), showing lagging-strand synthesis for the (+) reference strand.

### Mutations arising from *in vivo* oxidative stress

The *E. coli mutT* gene product is an 8-oxo-dGTP pyrophosphorylase [34] that efficiently removes the mutagenic product 8-oxo-dGTP from the cell [25]. In doing so, it powerfully prevents the occurrence of A·T→C·G transversions [25, 35, 36]. Some studies have indicated that the specific increase in A·T→C·G *in vivo* may be as high as 10,000-fold or more above the wild type level [25, 35, 36]. Here, we observe A·T→C·G at a frequency of 6.6 x 10^-7^ per base pair, a roughly 40-fold increase above the background. While this increase is much less than suggested by the *in vivo* values, this is an unavoidable consequence of the higher background of the duplex sequencing experiment, 2.1 x 10^-8^ (Table 1) versus 6 x 10^-10^ *in vivo* [25]. Nevertheless, it is important to note that our observed frequency of 6.6 x 10^-7^ per bp does compare reasonably to the frequency obtained for a series of A·T→C·G hotspot sites in the *lacI* gene (4 x 10^-6^) [25]. With regard to DNA sequence contexts (Table S6), our A·T→C·G mutations in *mutT* DNA show a preference for adenine as a 3’ template base with 190 out of 433 (44 %) occurring in 5’-**<u>A</u>**A-3’ contexts (**<u>A</u>** as template). This preference compares well to the context bias observed *in vivo*, where *mutT* spectra in the *lacI* gene revealed four strong hotspots at two 5’-T**<u>A</u>**A and two 5’-C**<u>A</u>**A sites [25].

### Perspective

The present work demonstrates the potential for using duplex sequencing in studies of bacterial mutagenesis. The ability to directly detect mutations in isolated DNA is a distinct advantage, as it avoids pitfalls of genetic selections and manipulations. A current disadvantage is that the background mutant frequency is relatively high, in the range of 10^-8^ to 10^-7^ per base pair, at least when compared to the very low *in vivo* values. The background is a likely consequence of DNA damage incurred during sample preparation and subsequent library construction. With improved insight into the sources of the DNA damage and into the *in vitro* mechanisms responsible for converting single-strand DNA damage to apparent duplex events, the background can presumably be further lowered significantly. Nevertheless, even at its current level very meaningful questions can already be addressed, as we have shown here for primary DNA replication errors and mutations resulting from *in vivo* oxidative stress. In the latter case, monitoring the presence of the mutagenic oxidation product 8-oxo-dGTP, producing diagnostic A·T→C·G events, is facilitated by the relatively low background of mutations at A·T base pairs. We envision application of duplex sequencing to other cases where (modest) mutator effects are observed or suspected, like those associated with certain stress conditions, starvations, and exposures to chemical agents and antibiotic treatments.

## Acknowledgements

This work was supported by the Intramural Research Program of the NIH, National Institute of Environmental Health Sciences [ZIA ES101905]. We acknowledge the help of the NIEHS Epigenomics and DNA Sequencing Core Facility for the sequencing runs, Dr. Brad Klemm and Dr. Libertad Garcia-Villada for their critical reading of the manuscript.

## References

[1] N. Ruiz, T.J. Silhavy, How Escherichia coli Became the Flagship Bacterium of Molecular Biology, J Bacteriol, (2022).

[2] J.W. Drake, B. Charlesworth, D. Charlesworth, J.F. Crow, Rates of spontaneous mutation, Genetics, 148 (1998) 1667–1686.

[3] R.M. Schaaper, Base Selection, Proofreading, and Mismatch Repair during DNA-Replication in Escherichia-Coli, J Biol Chem, 268 (1993) 23762–23765.

[4] J.W. Schroeder, P. Yeesin, L.A. Simmons, J.D. Wang, Sources of spontaneous mutagenesis in bacteria, Crit Rev Biochem Mol, 53 (2018) 29–48.

[5] R.M. Schaaper, B.N. Danforth, B.W. Glickman, Mechanisms of Spontaneous Mutagenesis - an Analysis of the Spectrum of Spontaneous Mutation in the Escherichia-Coli Laci Gene, J Mol Biol, 189 (1986) 273–284.

[6] S.J. Swerdlow, R.M. Schaaper, Mutagenesis in the lacI gene target of E-coli: Improved analysis for lacI(d) and lacO mutants, Mutat Res-Fund Mol M, 770 (2014) 79–84.

[7] H. Lee, E. Popodi, H. Tang, P.L. Foster, Rate and molecular spectrum of spontaneous mutations in the bacterium Escherichia coli as determined by whole-genome sequencing, Proc Natl Acad Sci U S A, 109 (2012) E2774–2783.

[8] H. Ochman, S. Elwyn, N.A. Moran, Calibrating bacterial evolution, P Natl Acad Sci USA, 96 (1999) 12638–12643.

[9] A.B. Williams, Spontaneous mutation rates come into focus in Escherichia coli, DNA Repair, 24 (2014) 73–79.

[10] M.W. Schmitt, S.R. Kennedy, J.J. Salk, E.J. Fox, J.B. Hiatt, L.A. Loeb, Detection of ultra-rare mutations by next-generation sequencing, P Natl Acad Sci USA, 109 (2012) 14508–14513.

[11] E.H. Ahn, K. Hirohata, B.F. Kohrn, E.J. Fox, C.C. Chang, L.A. Loeb, Detection of Ultra-Rare Mitochondrial Mutations in Breast Stem Cells by Duplex Sequencing, Plos One, 10 (2015).

[12] J.J. Salk, K. Loubet-Senear, E. Maritschnegg, C.C. Valentine, L.N. Williams, J.E. Higgins, R. Horvat, A. Vanderstichele, D. Nachmanson, K.T. Baker, M.J. Emond, E. Loter, M. Tretiakova, T. Soussi, L.A. Loeb, R. Zeillinger, P. Speiser, R.A. Risques, Ultra-Sensitive TP53 Sequencing for Cancer Detection Reveals Progressive Clonal Selection in Normal Tissue over a Century of Human Lifespan, Cell Rep, 28 (2019) 132–+.

[13] S.R. Kennedy, M.W. Schmitt, E.J. Fox, B.F. Kohrn, J.J. Salk, E.H. Ahn, M.J. Prindle, K.J. Kuong, J.C. Shen, R.A. Risques, L.A. Loeb, Detecting ultralow-frequency mutations by Duplex Sequencing (vol 9, pg 2586, 2014), Nat Protoc, 9 (2014) 2903–2903.

[14] N. Bhawsinghka, K.F. Glenn, R.M. Schaaper, Complete Genome Sequence of Escherichia coli BL21-AI, Microbiol Resour Ann, 9 (2020).

[15] H. Jeong, H.J. Kim, S.J. Lee, Complete Genome Sequence of Escherichia coli Strain BL21, Genome Announcements, 3 (2015).

[16] I.J. Fijalkowska, R.M. Schaaper, Effects of Escherichia-Coli Dnae Antimutator Alleles in a Proofreading-Deficient Mutd5 Strain, J Bacteriol, 177 (1995) 5979–5986.

[17] R.M. Schaaper, R.L. Dunn, The antimutator phenotype of E-coli mud is only apparent and results from delayed appearance of mutants, Mutat Res-Fund Mol M, 480 (2001) 71–75.

[18] K.H. Maslowska, K. Makiela-Dzbenska, J.Y. Mo, I.J. Fijalkowska, R.M. Schaaper, High-accuracy lagging-strand DNA replication mediated by DNA polymerase dissociation, Proc Natl Acad Sci U S A, 115 (2018) 4212–4217.

[19] H. Li, R. Durbin, Fast and accurate short read alignment with Burrows-Wheeler transform, Bioinformatics, 25 (2009) 1754–1760.

[20] Z.W. Lai, A. Markovets, M. Ahdesmaki, B. Chapman, O. Hofmann, R. McEwen, J. Johnson, B. Dougherty, J.C. Barrett, J.R. Dry, VarDict: a novel and versatile variant caller for next-generation sequencing in cancer research, Nucleic Acids Res, 44 (2016).

[21] A.R. Oller, R.M. Schaaper, Spontaneous Mutation in Escherichia-Coli Containing the Dnae911 DNA-Polymerase Antimutator Allele, Genetics, 138 (1994) 263–270.

[22] R.R. Iyer, A. Pluciennik, V. Burdett, P.L. Modrich, DNA mismatch repair: Functions and mechanisms, Chem Rev, 106 (2006) 302–323.

[23] R.M. Schaaper, R.L. Dunn, Spectra of spontaneous mutations in Escherichia coli strains defective in mismatch correction: the nature of in vivo DNA replication errors, Proc Natl Acad Sci U S A, 84 (1987) 6220–6224.

[24] L. Garibyan, T. Huang, M. Kim, E. Wolff, A. Nguyen, T. Nguyen, A. Diep, K.B. Hu, A. Iverson, H.J. Yang, J.H. Miller, Use of the rpoB gene to determine the specificity of base substitution mutations on the Escherichia coli chromosome, DNA Repair, 2 (2003) 593–608.

[25] R.G. Fowler, R.M. Schaaper, The role of the mutT gene of Escherichia coli in maintaining replication fidelity, Fems Microbiol Rev, 21 (1997) 43–54.

[26] R.M. Schaaper, R.L. Dunn, Escherichia-Coli Mutt Mutator Effect during Invitro DNA-Synthesis - Enhanced a-G Replicational Errors, J Biol Chem, 262 (1987) 16267–16270.

[27] P.L. Foster, H. Lee, E. Popodi, J.P. Townes, H. Tang, Determinants of spontaneous mutation in the bacterium Escherichia coli as revealed by whole-genome sequencing, Proc Natl Acad Sci U S A, 112 (2015) E5990–5999.

[28] B. Arbeithuber, K.D. Makova, I. Tiemann-Boege, Artifactual mutations resulting from DNA lesions limit detection levels in ultrasensitive sequencing applications, DNA Res, 23 (2016) 547–559.

[29] M. Costello, T.J. Pugh, T.J. Fennell, C. Stewart, L. Lichtenstein, J.C. Meldrim, J.L. Fostel, D.C. Friedrich, D. Perrin, D. Dionne, S. Kim, S.B. Gabriel, E.S. Lander, S. Fisher, G. Getz, Discovery and characterization of artifactual mutations in deep coverage targeted capture sequencing data due to oxidative DNA damage during sample preparation, Nucleic Acids Res, 41 (2013).

[30] K. Kino, M. Hirao-Suzuki, M. Morikawa, A. Sakaga, H. Miyazawa, Generation, repair and replication of guanine oxidation products, Genes Environ, 39 (2017).

[31] K. Kino, T. Kawada, M. Hirao-Suzuki, M. Morikawa, H. Miyazawa, Products of Oxidative Guanine Damage Form Base Pairs with Guanine, Int J Mol Sci, 21 (2020).

[32] M.L. Michaels, J.H. Miller, The Go System Protects Organisms from the Mutagenic Effect of the Spontaneous Lesion 8-Hydroxyguanine (7,8-Dihydro-8-Oxoguanine), J Bacteriol, 174 (1992) 6321–6325.

[33] I.J. Fijalkowska, P. Jonczyk, M.M. Tkaczyk, M. Bialoskorska, R.M. Schaaper, Unequal fidelity of leading strand and lagging strand DNA replication on the Escherichia coli chromosome, P Natl Acad Sci USA, 95 (1998) 10020–10025.

[34] S.K. Bhatnagar, M.J. Bessman, Studies on the Mutator Gene, Mutt of Escherichia-Coli - Molecular-Cloning of the Gene, Purification of the Gene-Product, and Identification of a Novel Nucleoside Triphosphatase, J Biol Chem, 263 (1988) 8953–8957.

[35] J.J. Vidmar, C.G. Cupples, Muty Repair Is Mutagenic in Mutt(-) Strains of Escherichia-Coli, Can J Microbiol, 39 (1993) 892–894.

[36] R.G. Fowler, S.J. White, C. Koyama, S.C. Moore, R.L. Dunn, R.M. Schaaper, Interactions among the Escherichia coli mutT, mutM, and mutY damage prevention pathways, DNA Repair, 2 (2003) 159–173.

