## Supplementary Tables S2-S6 for "Detection of DNA replication errors and 8-oxo-dGTP-mediated mutations in *E. coli* by Duplex DNA Sequencing"

**Table S1:** List of observed mutations by Duplex Sequencing (see separate excel sheet).

**Table S2:** Neighboring base sequence preferences for G·C→T·A substitutions in Duplex Sequencing (current data) in three different strains (significant biases are highlighted in red).

| G·C→T·A | | | |
| --- | --- | --- | --- |
|  | **Wild type** | ***mutL*** | ***mutT*** |
| 5’- A**G** | (31/160) = 19.4% | (7/33) = 21.2% | (27/166) = 16.3% |
| 5’- T**G** | (27/160) = 16.9% | (9/33) = 27.3% | (39/166) = 23.5% |
| 5’- G**G** | (50/160) = 31.3% | (10/33) = 30.3% | (51/166) = 30.7% |
| 5’- C**G** | (52/160) = 32.5% | (7/33) = 21.2% | (49/166) = 29.5% |
| **G**A-3’ | (30/160) = 18.8% | (8/33) = 24.2% | (56/166) = 33.7% |
| **G**T-3’ | (73/160) = 45.6% | (12/33) = 36.4% | (69/166) = 41.6% |
| **G**G-3’ | (39/160) = 24.4% | (6/33) = 18.2% | (24/166) = 14.5% |
| **G**C-3’ | (18/160) = 11.3% | (7/33) = 21.2% | (17/166) = 10.2% |

**Table S3:** Neighboring base sequence preferences for G·C→C·G substitutions in duplex sequencing (current data) for three different strains (significant biases are highlighted in red)

| G·C→C·G | | | |
| --- | --- | --- | --- |
|  | **Wild type** | ***mutL*** | ***mutT*** |
| 5’- A**G** | (14/136) = 10.3% | (3/30) = 10% | (18/97) = 18.6% |
| 5’- T**G** | (46/136) = 33.8% | (13/30) = 43.3% | (32/97) = 33% |
| 5’- G**G** | (35/136) = 25.7% | (3/30) = 10% | (18/97) = 18.6% |
| 5’- C**G** | (41/136) = 30.1% | (11/30) = 36.7% | (29/97) = 29.9% |
| **G**A -3’ | (64/136) = 47% | (12/30) = 40% | (32/97) = 33% |
| **G**T -3’ | (21/136) = 15.4% | (7/30) =2 3.3% | (16/97) = 16.5% |
| **G**G -3’ | (37/136) = 27.2% | (6/30) = 20% | (29/97) = 29.9% |
| **G**C -3’ | (14/136) = 10.3% | (5/30) = 16.7% | (20/97) = 20.6% |

**Table S4:** Neighboring-base sequence preferences for A·T→G·C substitutions in *mutL* DNA in Duplex Sequencing (current data) and *in vivo* studies (significant biases are highlighted in red). The lacI data (Maslowska *et al.* ) [1] represent the number of observed mutations at indicated sequence contexts divided by the number of mutationally detectable sites. The other two data (Duplex Sequencing and those of Lee *et al.* [2]) represent the number of mutations at the indicated sequence texts divided by the total number of observed A·T→G·C mutations.

| A·T→G·C | | | |
| --- | --- | --- | --- |
|  | **Duplex Sequencing (Present study)** | ***In vivo lacI* (Maslowska *et al*.)** | **Mutation accumulation**  **(Lee *et al*. )** |
| 5’- A**T** | (11/99) = 0.11 | (3/4) = 0.8 | (73/1141) = 0.06 |
| 5’- T**T** | (7/99) = 0.07 | (10/7) = 1.4 | (57/1141) = 0.05 |
| 5’- G**T** | (57/99) = 0.58 | (156/10) = 15.6 | (897/1141) = 0.79 |
| 5’- C**T** | (24/99) = 0.24 | (48/16) = 3 | (114/1141) = 0.10 |
| **T**A -3’ | (34/99) = 0.34 | (22/4) = 5.5 | (279/1141) = 0.24 |
| **T**T- 3’ | (19/99) = 0.19 | (9/3) = 3 | (277/1141) = 0.24 |
| **T**G -3’ | (20/99) = 0.20 | (57/18) = 3.2 | (281/1141) = 0.25 |
| **T**C -3’ | (26/99) = 0.26 | (47/12) = 3.9 | (304/1141) = 0.27 |

**Table S5:** Neighboring-base sequence preferences for G·C→A·T substitutions in *mutL* DNA in Duplex Sequencing (current data) and *in vivo* studies (significant biases are highlighted in red). The data are represented in a similar way as in Table S4.

| G·C→A·T | | | |
| --- | --- | --- | --- |
|  | **Duplex Sequencing (Present study)** | ***In vivo lacI* (Maslowska *et al*.)** | **Mutation accumulation**  **(Lee *et al.* )** |
| 5’- A**G** | (8/119) = 0.07 | (13/5) = 2.6 | (37/447) = 0.08 |
| 5’- T**G** | (33/119) = 0.28 | (151/31) = 4.9 | (79/447) = 0.18 |
| 5’- G**G** | (31/119) = 0.26 | (126/21) = 6.0 | (166/447) = 0.37 |
| 5’- C**G** | (47/119) = 0.39 | (52/13) = 4.0 | (165/447) = 0.37 |
| **G**A -3’ | (22/119) = 0.18 | (49/19) = 2.6 | (75/447) = 0.17 |
| **G**T -3’ | (19/119) = 0.16 | (59/16) = 3.7 | (56/447) = 0.13 |
| **G**G -3’ | (22/119) = 0.18 | (99/16) = 6.2 | (78/447) = 0.17 |
| **G**C -3’ | (56/119) = 0.47 | (146/21) = 7.0 | (238/447) = 0.53 |

**Table S6:** Neighboring base sequence preference of A·T→C·G substitutions in Duplex sequencing of *mutT* DNA (current data, significant biases are highlighted in red)

|  | ***mutT*** |
| --- | --- |
| 5’- A**A** | (108/433) = 0.25 |
| 5’- T**A** | (110/433) = 0.25 |
| 5’- G**A** | (120/433) = 0.28 |
| 5’- C**A** | (95/433) = 0.22 |
| **A**A - 3’ | (190/433) = 0.44 |
| **A**T- 3’ | (88/433) = 0.20 |
| **A**G - 3’ | (102/433) = 0.24 |
| **A**C - 3’ | (53/433) = 0.12 |

**References**

[1] K.H. Maslowska, K. Makiela-Dzbenska, J.Y. Mo, I.J. Fijalkowska, R.M. Schaaper, High-accuracy lagging-strand DNA replication mediated by DNA polymerase dissociation, Proc Natl Acad Sci U S A, 115 (2018) 4212-4217.

[2] H. Lee, E. Popodi, H. Tang, P.L. Foster, Rate and molecular spectrum of spontaneous mutations in the bacterium Escherichia coli as determined by whole-genome sequencing, Proc Natl Acad Sci U S A, 109 (2012) E2774-2783.
